# The C terminus of infectious bursal disease virus (IBDV) VP3 encodes a predicted intrinsically disordered region (IDR), which promotes the formation of cytoplasmic puncta and modulates their physical properties

**DOI:** 10.1101/2025.01.17.633518

**Authors:** A.J. Brodrick, A. J. Broadbent

## Abstract

The virus factories (VFs) of infectious bursal disease virus (IBDV) form through liquid-liquid phase separation (LLPS). A major component of the IBDV VF is the nonstructural protein VP3. Here, we predicted the full-length structure of the VP3 monomer and homodimer and employed molecular dynamics simulations to characterize their behavior. We identified the 36 amino acid carboxy(C)-terminus as a highly dynamic intrinsically disordered region (IDR). We then compared the cytoplasmic puncta that were made in the presence of the wild type (wt) VP3 with those made with a VP3 that lacked the C-terminus (VP3ΔC). Using live-cell imaging with fluorescent reporter tagged proteins, we found that VP3ΔC puncta were significantly less numerous (p<0.0001), smaller (p<0.0001), and more irregular in shape than puncta formed in the presence of wt VP3, demonstrating that the VP3 C terminal IDR promoted their formation. Moreover, by fluorescence recovery after photobleaching (FRAP), the VP3ΔC puncta had a significantly reduced mobile fraction (0.29) as compared to full-length VP3 puncta (0.70) (p<0.001), demonstrating that the VP3 C terminal IDR modulated their physical properties. However, the VP3ΔC puncta still exhibited liquid-like fusion events in the cytoplasm and were sensitive to treatment with aliphatic diols. Moreover, VP3 did not form puncta when expressed alone, and the removal of the C terminus did not abolish puncta formation completely. We propose that VP3 forms part of a higher order complex with other biomolecules to drive LLPS, and that the VP3 C terminal IDR modulates the physical properties of the resultant LLPS structures.

**Importance:** LLPS is a phenomenon of growing interest in cell biology. It is a part of the replication cycles of diverse viruses, but our understanding of the molecular basis that underpins the mechanism of phase separation is incomplete. We previously demonstrated that the birnavirus IBDV, a major agricultural pathogen, exploits LLPS in the formation of its VFs. Here, we have characterized the C-terminal 36 amino acid region of IBDV VP3 bioinformatically and by molecular dynamics simulations and found that it encodes a highly dynamic intrinsically disordered region (IDR). Furthermore, we found this region to promote the formation of cytoplasmic puncta and modulate their physical properties. This work contributes to a more detailed understanding of birnavirus replication at the molecular level, and to the study of LLPS as a phenomenon.

## Introduction

Viruses must regulate and organize biochemical processes to replicate efficiently, yet they also face strong selective pressures for gene and protein economy. One mechanism that achieves both of these goals is liquid-liquid phase separation (LLPS), where biomolecules spontaneously form liquid condensates in a liquid medium, separated from the surroundings by a phase transition barrier rather than the lipid membranes of typical organelles (1). LLPS is exploited by numerous viruses in the formation of their replicative compartments, including SARS-CoV-2 (2), adenoviruses (3), rabies viruses (4), Ebola viruses (5), measles virus (6), influenza viruses (7), and viruses with double stranded (ds)RNA genomes including reovirus (ReoV) (8), rotavirus (9), and birnaviruses (10).

LLPS is driven by the formation of molecular assemblages from biomolecules capable of multivalent interactions. It is distinguished from aggregation by the characteristically weak strength of the interactions, which permit diffusion of the biomolecules, while the cumulative effect of the interactions maintains the energetic favorability of phase separation (11, 12). Constituent components of biomolecular condensates formed through LLPS can be broadly categorized as “scaffold” or “client” biomolecules (13). Scaffold biomolecules are essential for phase separation to occur and contribute to the requisite multivalency that drives the phenomenon, whereas client biomolecules are recruited and phase separate, but would not do so in the absence of the scaffolds (13). The requirement of multivalent and weak interactions for phase separation means that intrinsically disordered proteins (IDPs) and proteins with intrinsically disordered regions (IDRs) are commonly found in LLPS structures, as these proteins and domains lack a single well-defined secondary structure, enabling them to participate in numerous weak or transient interactions as they explore their conformation space (12, 14). However, the requisite multivalency for phase separation can also be achieved through structured protein domains, meaning an IDPs or IDRs are not absolutely required for LLPS.

In ReoV, the non-structural protein µNS is considered to be a scaffold protein, which self-assembles into phase separated structures in the absence of other viral components (15), yet only the extreme N- and C-terminal sequences of µNS are likely to be disordered, and within the minimal region responsible for the formation of viral factory-like structures, there are two predicted alpha helices that have been proposed as principle interacting elements for µNS self-assembly (15). In contrast, in rotavirus, both NSP2 and NSP5 drive LLPS. NSP5 is an IDP that can assemble into several higher-ordered oligomers (16) and the Carboxy (C-)terminal region of NSP2 is flexible, allowing it to participate in domain-swapping interactions that are important in viroplasm formation (17). Despite these advances in our understanding of the molecular basis of phase separation in dsRNA viruses such as reoviruses and rotaviruses, comparatively little is understood regarding the mechanism underpinning the phase separation of the replicative compartments formed by birnaviruses.

Birnaviruses are nonenveloped, dsRNA viruses with a bipartite genome, and members of the *Birnaviridae* family include infectious bursal disease virus (IBDV) and infectious pancreatic necrosis virus (IPNV) which are pathogens of birds and fish, respectively, that are of economic importance to the poultry industry and aquaculture. Previously, we discovered that the viral replicative compartments of the *Birnaviridae* family, termed “virus factories” (VFs) also form through LLPS (10), but the molecular basis for phase separation of IBDV VFs has yet to be identified. A major component of the IBDV VF is viral protein (VP)3, which forms the matrix of the VF (10, 18). In this study we employed computational, cell culture, and biochemical approaches to better characterize the role IBDV VP3 plays in LLPS. We identified a predicted 36 residue C-terminal IDR in VP3 that we demonstrate promotes the formation of cytoplasmic puncta and modulates their physical properties, thus contributing to our knowledge of this virus, and to the field of LLPS.

## Results

### VP3 encodes a C-terminal domain with a predicted lack of secondary structure

IBDV encodes five proteins (VP1-5) across its two genome segments, with Segment A encoding a polyprotein that is cleaved to yield VP2,4 and 3 (Fig. 1A). Given that VP3 forms the matrix of the IBDV VF structures (19), we hypothesized that VP3 would act as a scaffold protein and show structural features conducive to driving LLPS. Only the structure of the VP3 “core” domains (residues 82-220) had previously been solved by X-ray crystallography (20), so we used AlphaFold2 to model the structure of the full-length IBDV VP3 monomer from IBDV strain PBG98 (Fig. 1B). The prediction was in good agreement with the crystal structure of the core domains, but notably lacked secondary structure in the C-terminal 36 residues (residues 221 – 257), hereafter referred to as the C-terminus. This correlated with a large predicted aligned error (PAE) in residue pairs in the C-terminus (Fig. S1A), suggesting a low confidence in the position of these residues relative to the rest of the structure (21). Moreover, these data were in agreement with the results of a disorder prediction algorithm (IUPred3 Short Disorder (22)), which indicated a propensity for increased disorder in the C-terminus as well as the N-terminus, and within two predicted “hinge” regions between structured portions of the core (Fig. 1C). This result inversely correlated with the Shannon entropy of the sequence (23)(Fig. S1B). Taken together, the residues of the C-terminus lacked predicted secondary structure, showed high IUPred3 disorder scores, and reduced sequence entropy, which are typical of a low-complexity IDR when measured by classical methods (24).

**Fig. 1.**
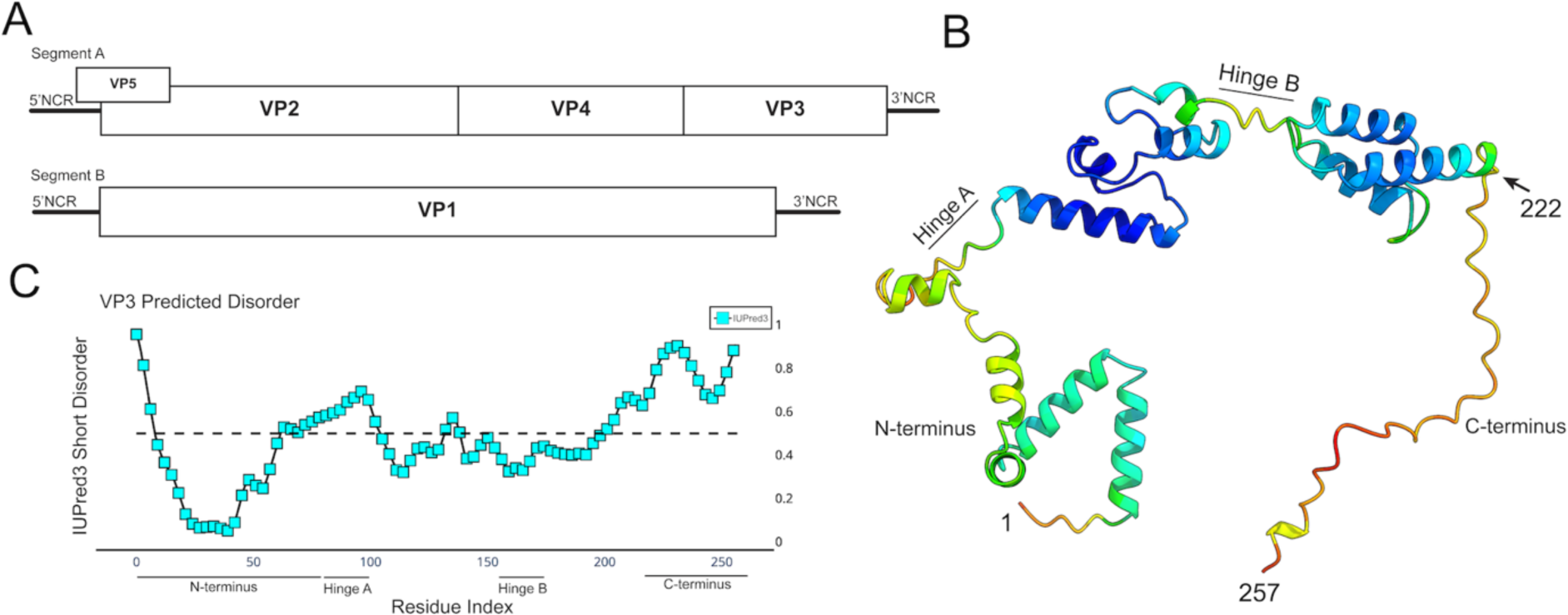
VP3 encodes a C-terminal domain with a predicted lack of secondary structure. A schematic of the IBDV genome architecture depicting Segment A encoding VP3 at the 3’ end of the coding strand, upstream of the 3’ non-coding region (NCR) and downstream of VP4 (A). A predicted structure for VP3, generated by AlphaFold2, visualized with UCSF ChimeraX, and colored by positional confidence (red indicates lower confidence; blue indicates higher confidence). The N terminus, C terminus and hinge regions are labelled (B). A disorder plot of the VP3 from IBDV strain PBG98, with IUPred3 short disorder plotted on the y axis against VP3 amino acid residue number on the x axis. The horizontal dashed line represents an IUPred3 Short Disorder of 0.5 and the N terminus, C terminus and hinge regions of VP3 are labelled (C).

### The VP3 C-terminal domain is predicted to be a highly dynamic intrinsically disordered region (IDR)

Molecular dynamics (MD) simulations of the VP3 monomer were employed to explore the predicted behavior of the C-terminus compared to VP3 as a whole. Each simulation was 20 nanoseconds in duration and the alpha carbons of VP3 were visualized as animated three-dimensional scatter plots of atomic co-ordinates (Movie S1). These data demonstrated that the C terminus explored a greater breadth of the conformation space than other domains of the VP3 monomer. To quantify the relative mobility of each residue, six independent simulations were run and the B factor, a metric associated with thermal motion, was calculated for each alpha carbon in each simulation. The average B factor values were plotted, which revealed an elevated B factor in the C-terminus (Fig. 2A), demonstrating its increased predicted mobility. However, B factor is an imperfect metric for distinguishing order from disorder, as ordered regions of proteins can undergo substantial thermal motion in MD simulations if they are adjacent to flexible “hinge” or “loop” regions (25). Therefore, to determine if the behavior of the C-terminus differed from other regions of the protein, we conducted a pairwise correlation between the movement of the atoms across the VP3 monomer (atomic motion correlation analysis) and plotted the results as a heatmap (Fig. 2B). This revealed that the movement of the C-terminus was not well correlated with other regions of the protein (Fig. 2B, bottom right corner), suggesting it moves independently of the rest of the protein. As the C terminus is 36 amino acids in length, next we determined the atomic motion of equally-sized sections of 36 amino acids in length across the whole VP3 protein and compared these data to the C terminus, to determine whether a given region was significantly more ordered than the C-terminus or not (Fig. 2C). If a given region of the protein was significantly more ordered than the C-terminus, it had a p value below the 0.05 cut-off (red line), whereas if a particular region was not significantly more ordered than the C-terminus, then the p value was higher and hence gave a peak (Fig. 2C). The VP3 protein was significantly more ordered than the C-terminus at all regions except the C-terminus itself, and amino acids 74-96 and 149-163, both of which correspond to the predicted unstructured hinge regions of the protein. Taken together with the calculated B-factor values and the broad conformational landscape explored by the C-terminus, the results of MD simulations demonstrated that the VP3 C-terminus is predicted to be a highly dynamic IDR.

**Fig. 2.**
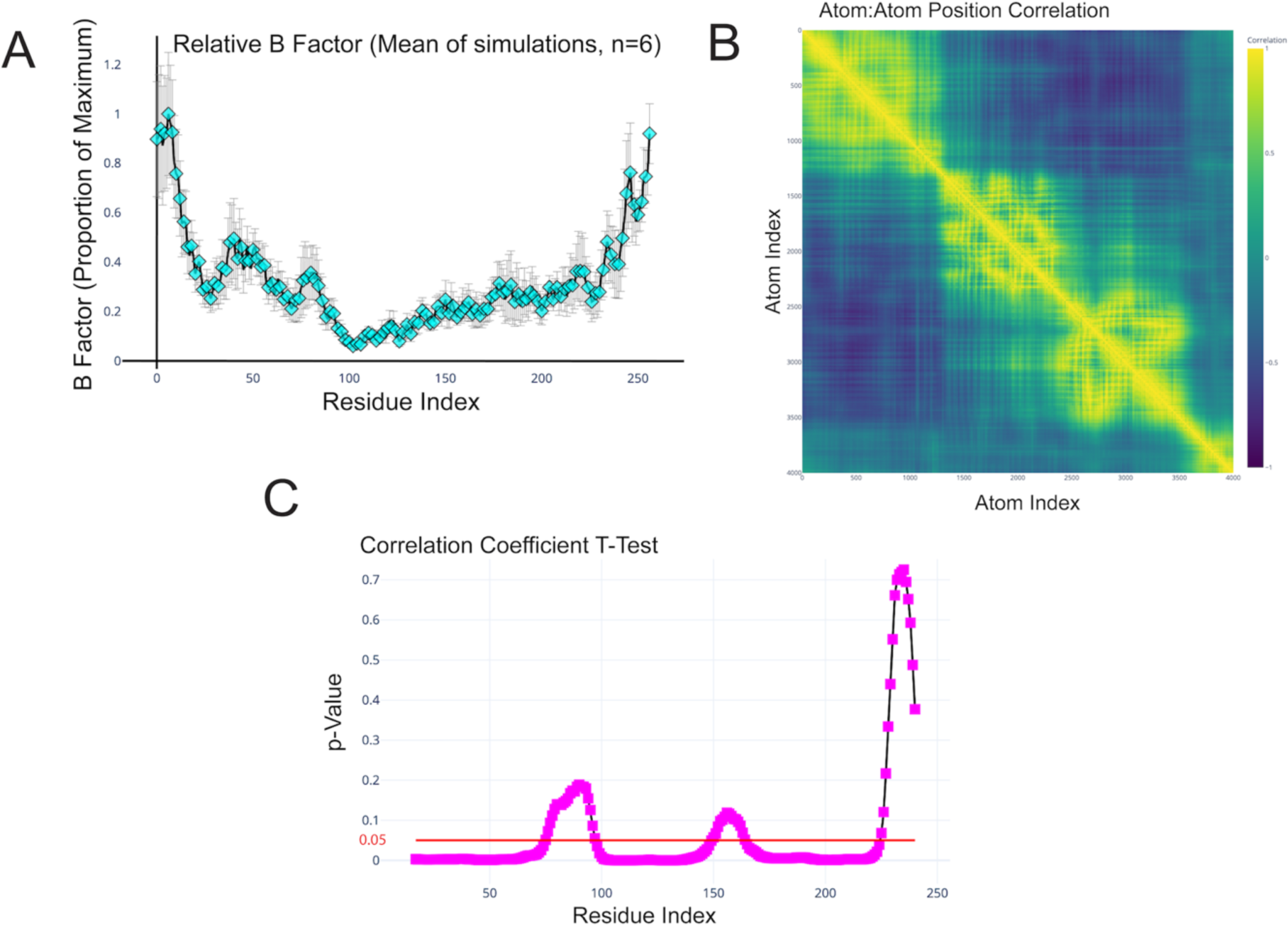
The VP3 C-terminal domain is predicted to be a highly dynamic intrinsically disordered region (IDR). The mean relative B-factor, calculated from MD simulations of the VP3 monomer (6 independent 20 nanosecond simulations), was plotted on the y axis against VP3 amino acid residue number on the x axis. Error bars represent standard error of the mean (SEM) (A). The Pearson’s product moment correlation coefficient between the trajectories of each pair of atoms was calculated and plotted as a heatmap. Yellow represents perfect positive correlation, purple represents perfect negative correlation, blue/teal suggests a lack of correlation (B). The movement of the atoms (average positional correlation coefficient) within the 36 amino acids of the C terminus were compared to equally-sized windows of 36 amino acids in length spanning the whole length of the VP3 protein sequence, and the p-values were plotted on the y axis against the VP3 amino acid residue number on the x axis. Regions of the VP3 region that were significantly more ordered than the C-terminus had a p value below the 0.05 cut-off (red line) (C).

### IBDV VP3 can exist as a homodimer with two C terminal dynamic IDRs extending from the core

In infected cells, VP3 can exist as a homodimer with the dimerization domain of VP3 predicted to lie within the N-terminus of the protein (26). We therefore generated a predicted structure for the VP3 homodimer using AlphaFold3. The VP3 dimer model featured interactions between the N-termini of the monomers, but by virtue of the flexible hinge regions predicted in the monomer, the “core” regions wrapped around the N-termini, leading to an overall compact conformation. The C-termini of both subunits extended from the compact cores and exhibited the same low-confidence we noted in the monomer model (Fig. 3 A and B). MD simulations of the VP3 homodimer revealed the C-termini of both subunits remained highly flexible and dynamic (Movie S2). The resultant trajectories of the alpha carbons in the VP3 homodimer were visualized as an ensemble of ten evenly-spaced timepoints from a single 20 nanosecond molecular dynamics simulation (Fig. 3C), demonstrating the breadth of the conformation space explored by the C termini. B-factors were calculated for the alpha carbons in the homodimer, and these data indicated that the elevated C-terminal B-factors observed in the monomer simulations were also observed for both subunits of the dimer (Fig. 3D).

**Fig. 3.**
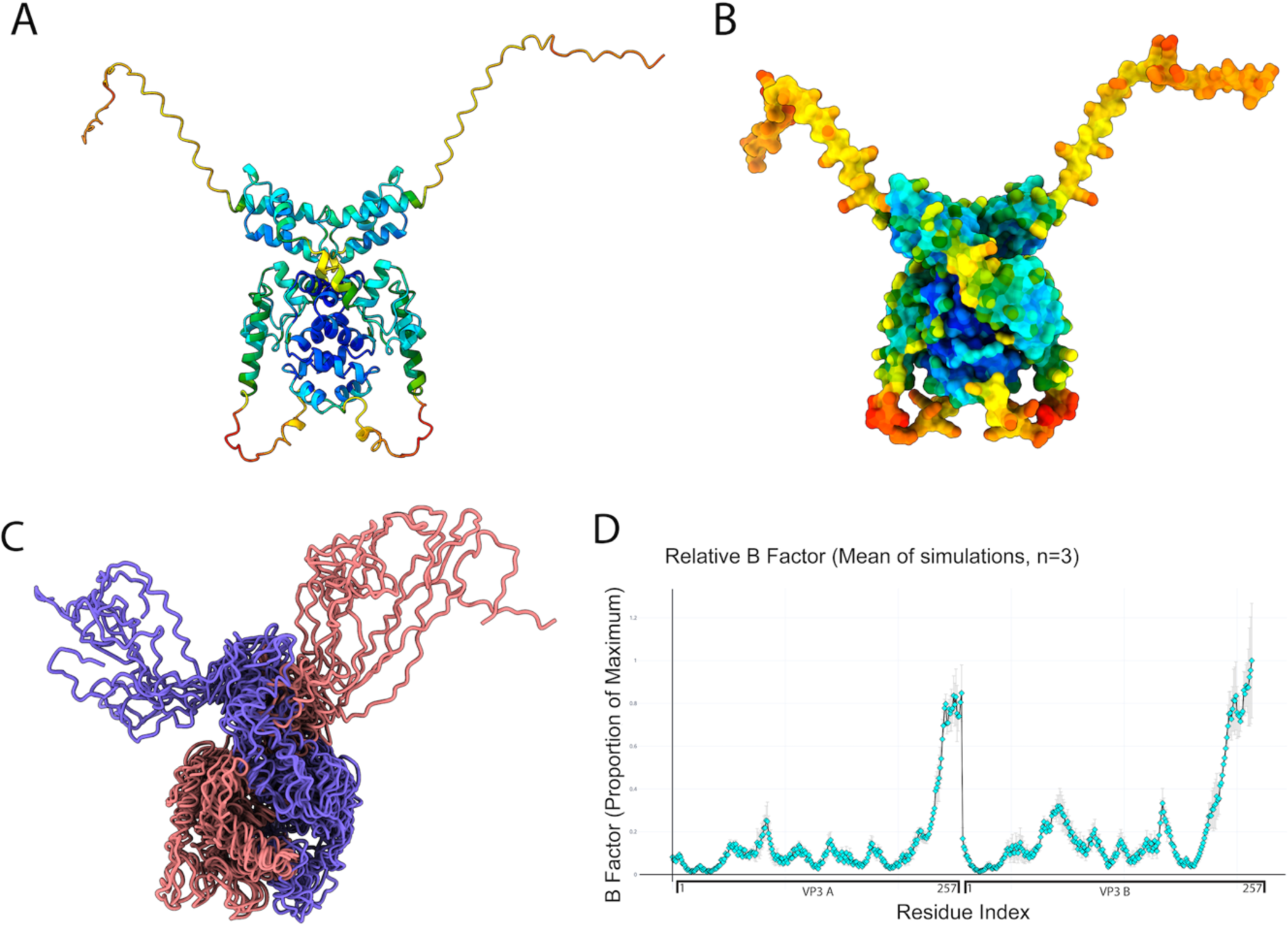
IBDV VP3 can exist as a homodimer with two C terminal dynamic IDRs extending from the core. A cartoon rendering of the VP3 homodimer as predicted by AlphaFold3, visualized in UCSF ChimeraX, and colored by positional confidence (red indicates lower confidence; blue indicates higher confidence) (A). A solvent-accessible surface of the VP3 homodimer, visualized by UCSF ChimeraX, and colored by positional confidence (red indicates lower confidence; blue indicates higher confidence) (B). An ensemble of structures representing ten evenly-spaced timepoints of a single 20 nanosecond molecular dynamics simulation. Structures were extracted from the trajectory and rendered as an ensemble in UCSF ChimeraX without alignment, colored by monomer (one monomer in purple, the other in pink) (C). The mean relative B-factor was calculated from the molecular dynamics simulations of the VP3 homodimer (6 independent 20 nanosecond simulations) and plotted on the y axis against the amino acid residue number of two VP3 monomers (A and B) concatenated on the x axis. Error bars represent SEM (D).

### The C terminal IDR is not required for the recruitment of IBDV VP3 into VFs

To determine the behavior of VP3 in cells, it was first necessary to develop a reporter-tagged fusion construct to permit live cell imaging experiments. We therefore prepared a panel of VP3 proteins fused to different reporters. All tags were fused to the N-terminus of VP3, as previous reports revealed that C-terminal fusions prevented the rescue of recombinant viruses by reverse genetics and might therefore interrupt the function(s) of the VP3 C-terminus (27). Reporters included tetracysteine (TC), enhanced green fluorescence protein (eGFP), and mNeonGreen (mNG) (28) (Fig. S2). Cells transfected with TC::VP3 were chemically labeled with FLaSH prior to imaging, and cells transfected with an untagged VP3 control were analyzed by immunofluorescence microscopy (IFM) with a mouse monoclonal antibody raised against full-length VP3 (*α*-VP3) (29). Interestingly, when expressed alone, untagged VP3 assumed a predominantly diffuse or moderately reticulated cytoplasmic staining pattern in transfected DF-1 cells, and the pattern of the TC::VP3 and mNG::VP3 fluorescence signals were also consistent with this observation. In contrast, eGFP::VP3 formed large aggregates in the cytoplasm of transfected cells, and so subsequent experiments were not performed with this reporter. When we simultaneously co-transfected DF-1 cells with the reporter::VP3 plasmids and infected them with IBDV strain PBG98, we observed a redistribution of the fluorescence signal into cytoplasmic puncta. These data demonstrate that when expressed alone, VP3 did not spontaneously phase separate into cytoplasmic puncta, however, when expressed in infected cells, VP3 was recruited to the VFs. We selected mNG as a reporter for use in subsequent experiments due to its nature as a constitutively fluorescent protein, functionality in live cells, known low propensity for multimerization (15, 28), and performance in our screen.

We next aimed to compare the behavior of the full-length (wild type (wt)) VP3 with a VP3 molecule that lacked the 36 residue C-terminus. We therefore truncated the mNG::VP3 construct after residue 221 of VP3, to form mNG::VP3ΔC. An alignment of the predicted structures of VP3 and VP3ΔC using Alphafold3 and ChimeraX revealed an identical secondary structure configuration, with differences between the structures contributed only by the flexible linker regions, demonstrating that there were no predicted folding defects in the remaining protein (Fig. 4A). We then transfected DF-1 cells with the mNG::VP3ΔC construct, and stained the cells with *α*-VP3, revealing a complete overlap of the mNG and VP3 signals, and a cytoplasmic distribution matching that of full-length VP3. Moreover, colocalization analysis of the mNG::VP3ΔC signal and the anti-VP3 signal revealed a Manders’ coefficient of 0.903 and Spearman’s rank correlation of 0.793 (Fig. S3), demonstrating that truncating the C terminus did not substantially affect the intracellular localization of the protein in transfected cells, or the binding of the anti-VP3 antibody. To determine if deletion of the C-terminus altered the recruitment of exogenous VP3 to VFs, DF-1 cells were either transfected with either mNG::VP3 or mNG::VP3ΔC alone, or simultaneously infected with IBDV strain PBG98 and transfected with either mNG::VP3 or mNG::VP3ΔC (Fig. 4B). The mNG signal was observed to be redistributed from diffusely cytoplasmic to punctate in infected cells transfected with either mNG::VP3 or mNG::VP3ΔC, indicating that deletion of the C-terminus did not prevent the recruitment of exogenous VP3 to IBDV VFs.

**Fig. 4.**
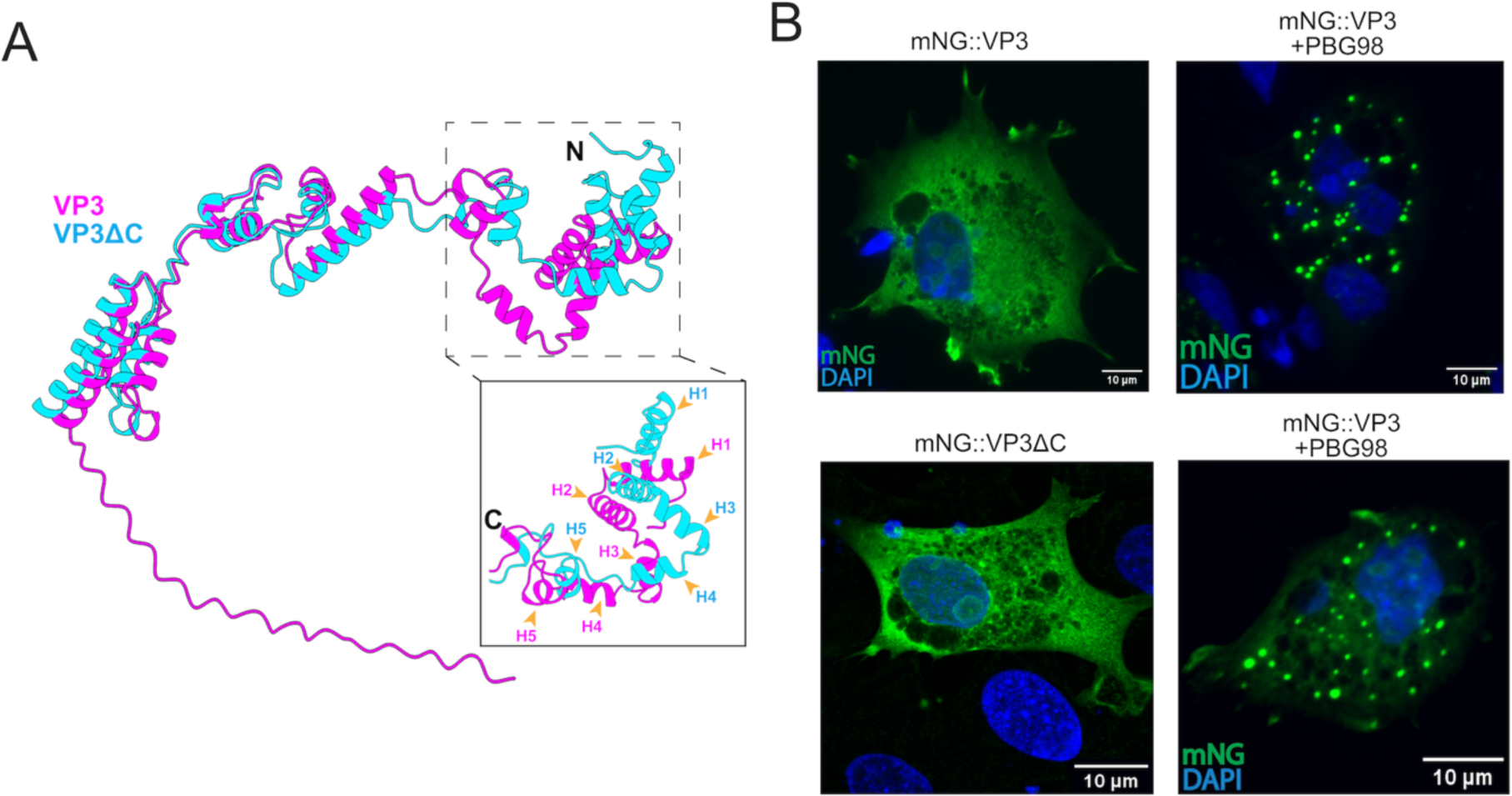
The C terminal IDR is not required for the recruitment of IBDV VP3 into VFs. AlphaFold3 predictions for the wt VP3 (magenta) and the VP3ΔC (cyan) were aligned with ChimeraX and overlaid (N terminal helices numbered H1-5) (A). DF-1 cells were transfected with mNG::VP3 alone, or were transfected with mNG::VP3 and simultaneously infected with IBDV strain PBG98 (MOI 5) (B). DF-1 cells were transfected with mNG::VP3ΔC alone, or transfected with mNG::VP3ΔC and simultaneously infected with strain IBDV strain PBG98 (MOI 5) (C). Cells were fixed at 12hpi, and the nuclei were stained with DAPI (blue) and the mNG signal is shown in green. Scale bars, 10µm.

### The IBDV VP3 C terminal IDR is not required for the formation of cytoplasmic puncta

Although our data indicated that VP3 recruitment to IBDV VFs was independent of the C-terminus, the VFs formed by co-infection and transfection contained a mixture of exogenously expressed VP3ΔC and full-length VP3 derived from the infecting virus, which prevented an analysis of how the VP3 C terminus affected the formation and properties of the VFs. To address this, we first aimed to rescue a molecular clone of IBDV with a VP3 that lacked the C terminus. We have an in-house reverse genetics system that involves two plasmids, one encoding Segment A (pIBDV-SegA), and one encoding Segment B (pIBDV-SegB) (30). As VP3 is located at the C terminus of the polyprotein that is translated from Segment A (Fig. 1A), it was possible to remove the C terminus of VP3 by truncating the Segment A coding region by 36 amino acids, to form pIBDV-SegAΔC. Unfortunately, this virus failed to rescue, but we reasoned that we could make use of the reverse genetics plasmids to study the IBDV puncta in the absence of infection. Given that VP3 failed to form cytoplasmic puncta when expressed alone, we hypothesized that it required the interaction with other viral proteins, and/or RNA, to form VFs. We hypothesized that simultaneous transfection with both wt reverse genetics plasmids would induce the punctate phenotype, as together they encode all viral proteins, and permit the synthesis of viral RNA (vRNA) and the double-stranded RNA (dsRNA) genome, which is known to localize to VFs (18). We therefore transfected DF-1 cells with the Segment A reverse genetics plasmid and visualized expression of VP3 by IFM, and we transfected DF-1 cells with a Segment B reverse genetics plasmid that we modified to express an HA-tag as we did not have an antibody that recognized the Segment B product, VP1 (the HA tag was inserted between the 5’ untranslated region (UTR) and the VP1 coding sequence (pIBDV-HA::SegB)). Both Segment A and B, when expressed alone, revealed a diffuse cytoplasmic staining pattern (Fig. 5A), demonstrating that neither segment expressed independently recapitulated cytoplasmic puncta. In contrast, when we transfected DF-1 cells with a mixture of plasmids encoding Segments A and B, and visualized the resultant VP3 distribution by IFM, we observed the appearance of cells with punctate cytoplasmic inclusions, which indicated that when simultaneously expressed, the IBDV reverse genetics system was capable of recapitulating the punctate phenotype (Fig. 5A), consistent with VP3 requiring additional factors to induce cytoplasmic puncta. Next, we transfected DF-1 cells with reverse genetics plasmids pIBDV-SegAΔC and pIBDV-HA::SegB (Fig. 5B). Similarly to the wt plasmids, the pIBDV-SegAΔC and pIBDV-HA::SegB took on a diffuse cytoplasmic staining pattern when expressed alone, however, when cells were co-transfected with a mixture of pIBDV-SegAΔC and pIBDV-SegB, we detected cytoplasmic puncta (Fig. 5B), demonstrating that the VP3 C-terminal IDR was not an absolute requirement for the appearance of cytoplasmic puncta.

**Fig. 5.**
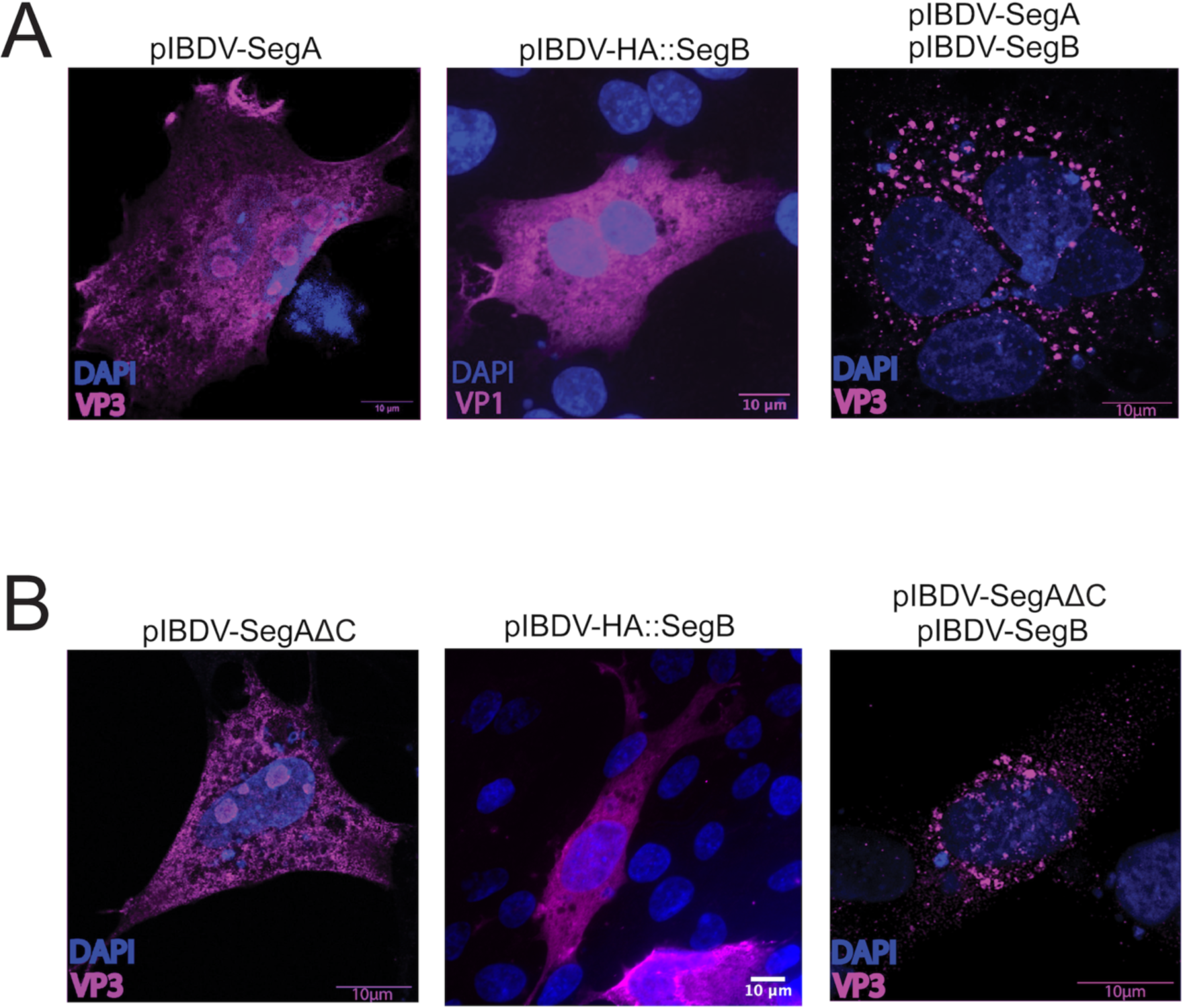
The IBDV VP3 C terminal IDR is not required for the formation of cytoplasmic puncta. DF-1 cells were either transfected with pIBDV-SegA or pIBDV-HA::SegB, and fixed and stained with anti-VP3 or anti-HA antibodies at 18 hpt, or DF-1 cells were simultaneously transfected with both pIBDV-SegA and pIBDV-SegB, and fixed and stained with an anti-VP3 antibody at 18 hpt (A). DF-1 cells were either transfected with pIBDV-SegAΔC or pIBDV-HA::SegB, and were fixed and stained with anti-VP3 or anti-HA antibodies at 18 hpt, or DF-1 cells were simultaneously transfected with both pIBDV-SegAΔC and pIBDV-SegB, and fixed and stained with an anti-VP3 antibody at 18 hpt. The nuclei were stained with DAPI (blue) and the VP3 or HA-VP1 signals are shown in magenta. Scale bars, 10µm (B).

### The IBDV VP3 C terminal IDR promotes the formation of cytoplasmic puncta and influences their size and shape

The requirement for fixing, staining, and IFM precluded live cell imaging, and therefore it was not possible to assess whether the VP3 IDR affected the physical properties of the puncta using the system we had developed. To address this deficit, we modified our pIBDV-SegB plasmid by adding mNG to the C-terminus of the VP1 coding sequence (pIBDV-SegB::mNG) (Fig. S4). We then transfected cells with either a mixture of pIBDV-SegA and pIBDV-SegB::mNG, or a mixture of pIBDV-SegAΔC and pIBDV-SegB::mNG, and we detected the presence of mNG-positive cytoplasmic puncta that colocalized with the VP3 signal (Fig. S4). This system was therefore capable of generating reporter-tagged puncta that could be imaged live. Next, fifty randomly selected cells from each of three independent replicates of cells transfected with either pIBDV-SegA and pIBDV-SegB::mNG, or pIBDV-SegAΔC and pIBDVSegB::mNG were imaged, and the puncta were counted. Although cells containing puncta were observed in the absence of the C-terminus, they were detected at a significantly reduced rate (p<0.0001) (Fig. 6A). Furthermore, among the cells that did contain puncta, the mean number of puncta per cell was significantly reduced in the ΔC system compared to wt (p<0.0001) (Fig 6B). Taken together, these data demonstrate that although the IBDV VP3 C terminal IDR was not required for the formation of cytoplasmic puncta, it did promote their formation.

**Fig. 6.**
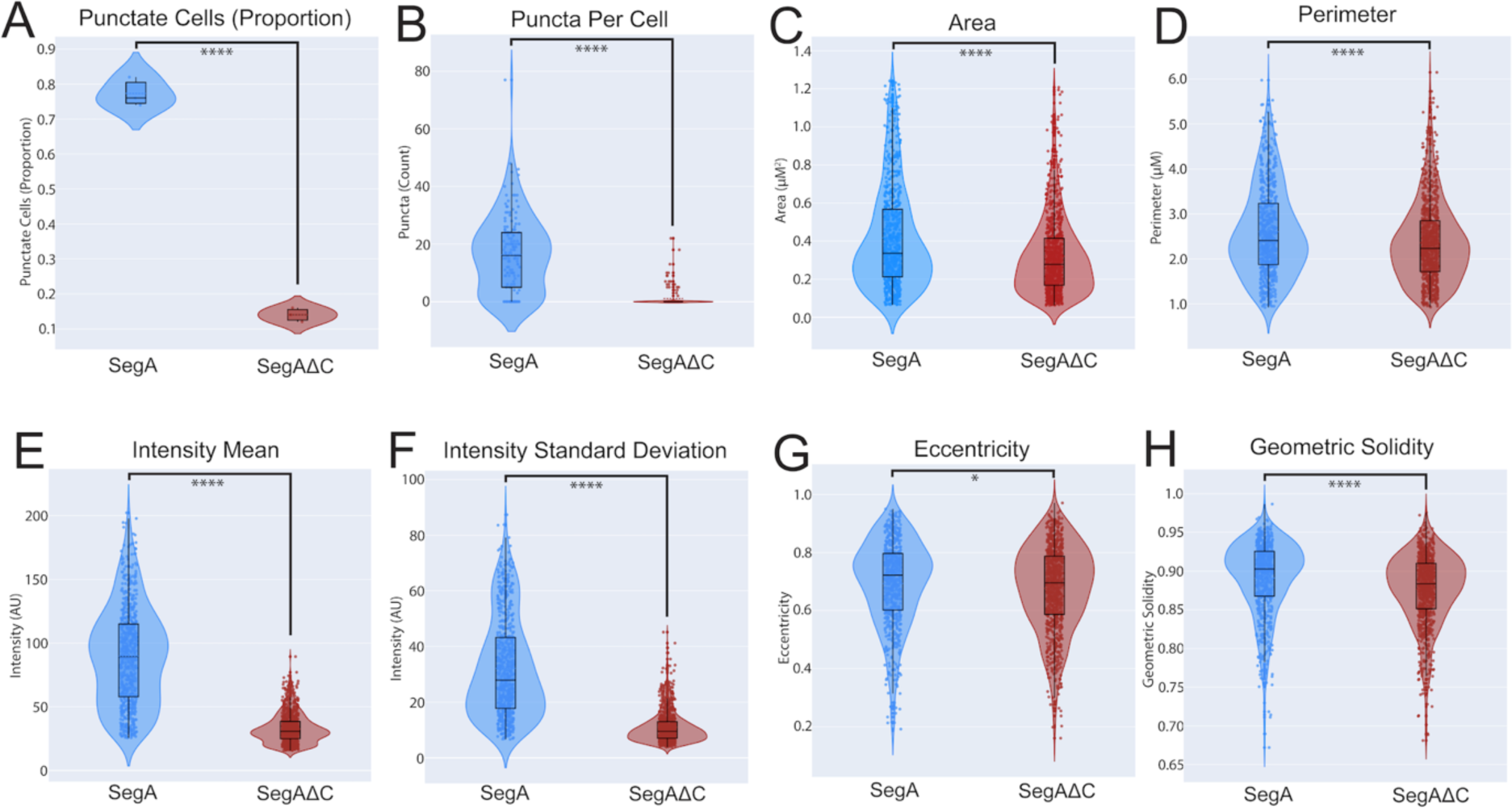
The IBDV VP3 C terminal IDR promotes the formation of cytoplasmic puncta and influences their size and morphology. DF-1 cells were co-transfected with either pIBDV-SegA and pIBDV-SegB::mNG, or pIBDV-SegAΔC and pIBDV-SegB::mNG. From each of 3 replicates, 50 cells were randomly selected at 34hpt, and their phenotype (punctate or non-punctate) was classified (A), and the puncta per cell were counted (B). Fifty cells prepared in the same manner were then imaged individually by confocal microscopy, and puncta were segmented by watershed segmentation and the area, perimeter, mean fluorescence intensity, intensity standard deviation, eccentricity, and geometric solidity (a measure of morphological “roundness” or “smoothness”) were measured with the Scikit-image python module (C - H).

Six morphological properties of the puncta were then measured: area, Crofton perimeter, single-puncta mean fluorescence intensity, single-puncta intensity standard deviation, eccentricity, and geometric solidity (a measure of morphological “roundness” or “smoothness”). Puncta that formed in the presence of wt VP3 had a significantly larger average area and perimeter than puncta formed in the presence of VP3ΔC (p<0.0001) (Fig. 6C and D). Moreover wt VP3 puncta had a significantly greater mean fluorescence intensity, and a greater single-puncta intensity variability than VP3ΔC puncta (p<0.0001) (Fig. 6E and F), consistent with the VP3 C terminal IDR contributing to the size and intensity of the puncta. Finally, wt VP3 puncta had a significantly higher mean eccentricity than VP3ΔC puncta (p<0.05), and exhibited significantly higher mean geometric solidity (p<0.0001), indicating that the VP3 C terminal IDR contributed to the “roundness” or “smoothness” of the puncta. Taken together, these data demonstrate that the VP3 C terminal IDR influenced the size and shape of the cytoplasmic puncta.

### The VP3 C-terminus affects the physical properties of VFs

Finally, we aimed to characterize differences in the physiochemical properties of puncta formed in the presence of the wt VP3 or the VP3ΔC. Classification of structures as being LLPS typically involves several assays for characterizing liquid-like properties, including common morphological traits, dynamic behavior, disruption upon treatment with small aliphatic diols, and rapid internal diffusion (31). LLPS structures are typically dynamic, and exhibit mobility within their cellular compartment, and may undergo fission (the splitting of a single structure into two or more) and fusion (the merging of two or more structures into one upon collision).

To assess differences in the physical properties of the puncta quantitatively, we performed fluorescence recovery after photobleaching (FRAP), a technique which characterizes the diffusion of molecules within a structure. Briefly, a small region of the fluorescently labeled puncta was photobleached by a laser pulse, permanently inactivating the fluorophores in that region, and the mean fluorescence of the bleached region was then measured over time. Brownian motion of the constituent molecules caused the inactivated fluorophores to diffuse away from the bleach site and unbleached molecules from outside the bleach site to diffuse in, resulting in a net recovery of fluorescence, and a curve was fitted to these data points, from which the “liquidity” of the structure could be described. Briefly, we transfected DF-1 cells with either a mixture of pIBDV-SegA and pIBDV-SegB::mNG (Fig. 7A), or a mixture of pIBDV-SegAΔC and pIBDV-SegB::mNG (Fig 7B) and we captured time lapse series by live confocal microscopy at 14 hpt, with a single iteration of point bleaching on the 10^th^ frame of each series. Intensity data was extracted from the raw image series with ImageJ, and a custom Python script implementing the EasyFrap Web algorithm (32) was used to calculate recovery curves. We observed good agreement between independent replicates for each condition, and a notable difference between the recovery behavior of puncta formed in the presence of wt VP3 and the VP3ΔC (Fig. 7). Puncta in cells expressing VP3ΔC exhibited a partial recovery in fluorescence after photobleaching, but the mobile fraction was calculated as only 0.291, indicating that only 29.1% of the molecules exhibited any diffusion. This was in sharp contrast to the behavior of puncta formed in cells expressing the wt VP3, where although the recovery half-time was longer at 3.517 seconds (compared to 0.341 seconds in the VP3ΔC puncta), the calculated mobile fraction was 0.704, indicating that 70.4% of the molecules exhibited diffusion, a significantly greater proportion compared to the VP3ΔC puncta (p<0.01). These data demonstrate that the VP3 C-terminal IDR affected the physical properties of the VFs.

**Fig. 7.**
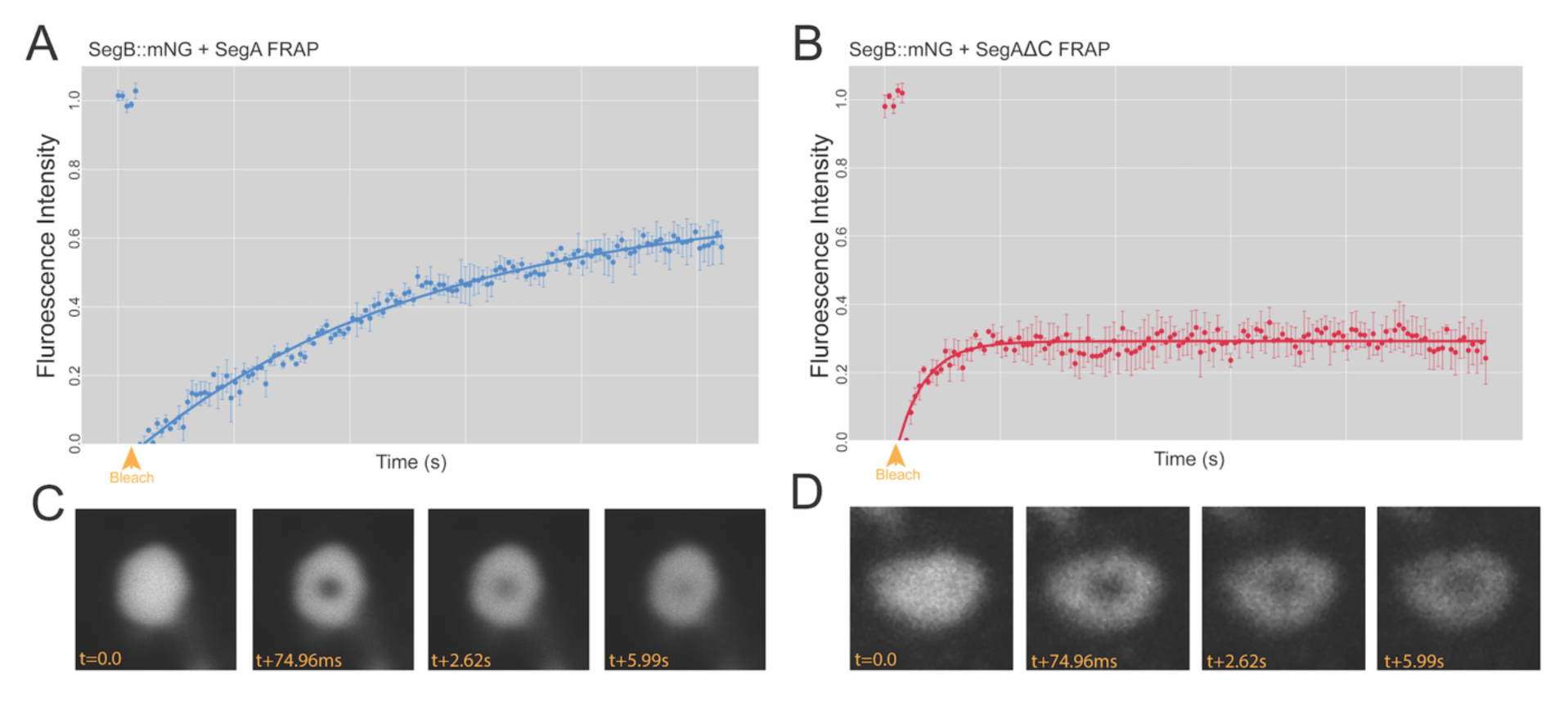
The VP3 C terminal IDR significantly contributes to the mobile fraction of the moleculaes within the cytoplasmic puncta. DF-1 cells were co-transfected with a mixture of pIBDV-SegA and pIBDV-SegB::mNG and a time lapse series was captured by live confocal microscopy at 14 hpt with a single iteration of point bleaching on the 10^th^ frame (yellow arrowhead). Intensity data was extracted and used to calculate a FRAP recovery curve. Results are from 3 replicates; error bars = SEM (A). DF-1 cells were co-transfected with a mixture of pIBDV-SegAΔC and pIBDV-SegB::mNG and a FRAP recovery curve generated in the same way (B). Representative recovery images for a photobleached puncta formed in the presence of pIBDV-SegA with the time post bleach (t) shown in milliseconds (ms) (C). Representative recovery images for a photobleached puncta formed in the presence of pIBDV-SegAΔC with the time post bleach shown in ms (D).

To assess other behaviors associated with LLPS, we transfected DF-1 cells with either a mixture of pIBDV-SegA and pIBDV-SegB::mNG (Fig. 8A), or a mixture of pIBDV-SegAΔC and pIBDV-SegB::mNG (Fig. 8B), and observed the cells by live-cell confocal microscopy, capturing images every 5 frames to make a time lapse. Interestingly, we observed that both the cytoplasmic puncta made in the presence of either wt VP3 or VP3ΔC were mobile, moving throughout the cytoplasm on similar, rapid timescales, and underwent fusion events (Fig. 8), indicating that these key liquid-like properties were retained in the absence of the VP3 C-terminus. Furthermore, we characterized the response of the puncta to treatment with small aliphatic diols, which have been observed to disrupt LLPS structures (31). Briefly, we transfected DF-1 cells with either a mixture of pIBDV-SegA and pIBDV-SegB::mNG (Fig. 9A), or a mixture of pIBDV-SegAΔC and pIBDV-SegB::mNG (Fig. 9B), and treated the transfected cells with 1,3-Propanediol (1,3-PD), an aliphatic diol that exhibits lower toxicity than the more commonly used 1,6-Hexanediol (1,6-HD), while capturing time lapse images (9). Although the toxicity of 1,3-PD was reduced as compared to 1,6-HD, eventually the cells did succumb to the treatment, meaning our observations were limited to approximately ten minutes post-treatment where we could reliably image the puncta. Over the course of the time lapse, 1,3-PD caused a moderate degree of disruption to puncta made with both wt VP3 or VP3ΔC (Fig 9). The margins of most puncta exhibited a loss of definition, although complete dissolution was only observed for a subset of smaller puncta. Both puncta formed in the presence of wt VP3 and in the presence of VP3ΔC were sensitive to diol treatment, but the responses varied too widely between cells and between puncta within a single cell to quantitatively distinguish differences in the behavior of the puncta made with either wt VP3 or VP3ΔC.

**Fig. 8.**
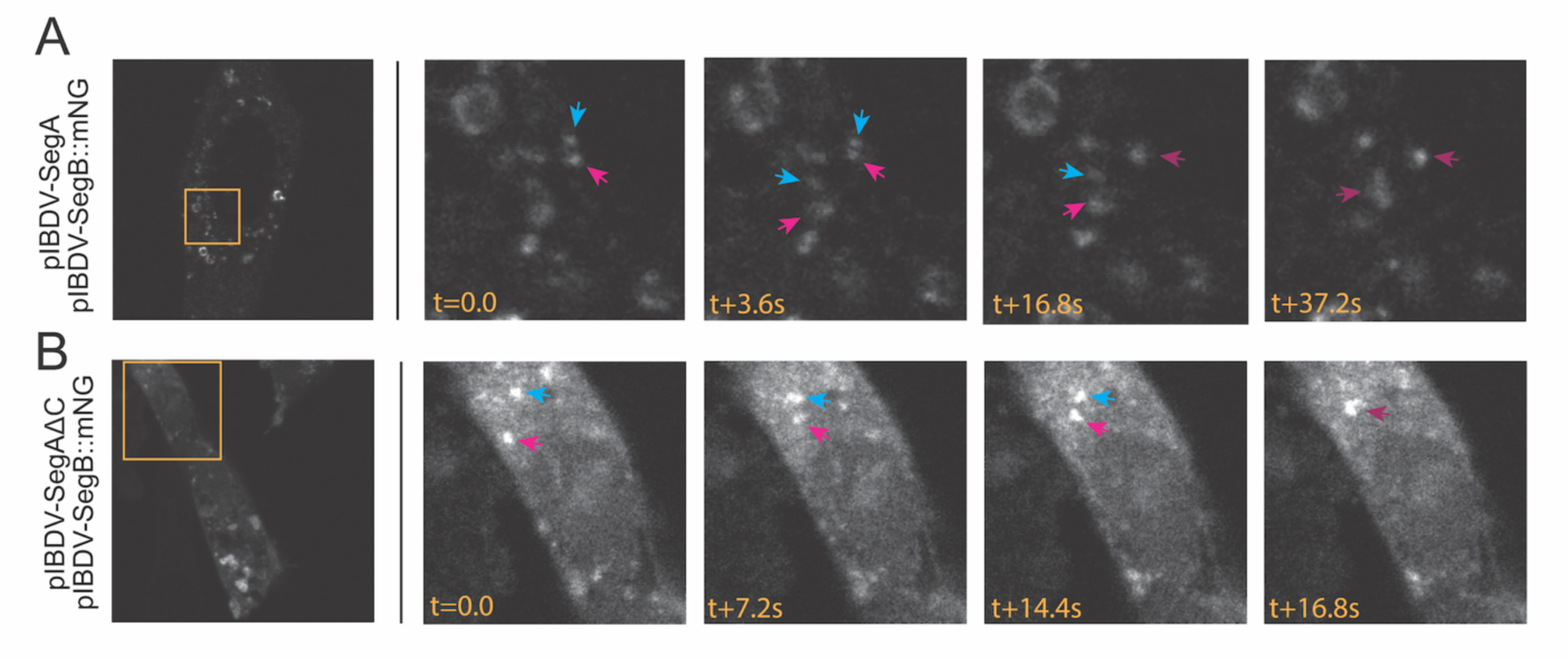
Puncta formed in the presence or absence of the VP3 C terminal IDR are dynamic. DF-1 cells were transfected with a mixture of pIBDV-SegA and pIBDV-SegB::mNG (A), or DF-1 cells were transfected with a mixture of pIBDV-SegAΔC and pIBDV-SegB::mNG (B). At 34hpt, cells were imaged by live-cell confocal microscopy, capturing images every 5 frames to make a time lapse. Still frames from the timelapses are shown. The leftmost images show whole-cell images (mNG signal shown in white), with stills from the inset regions (orange boxed regions) shown thereafter. The timestamp (t) of imaging is shown in seconds (s). Arrows show individual puncta that undergo fusion.

**Fig. 9.**
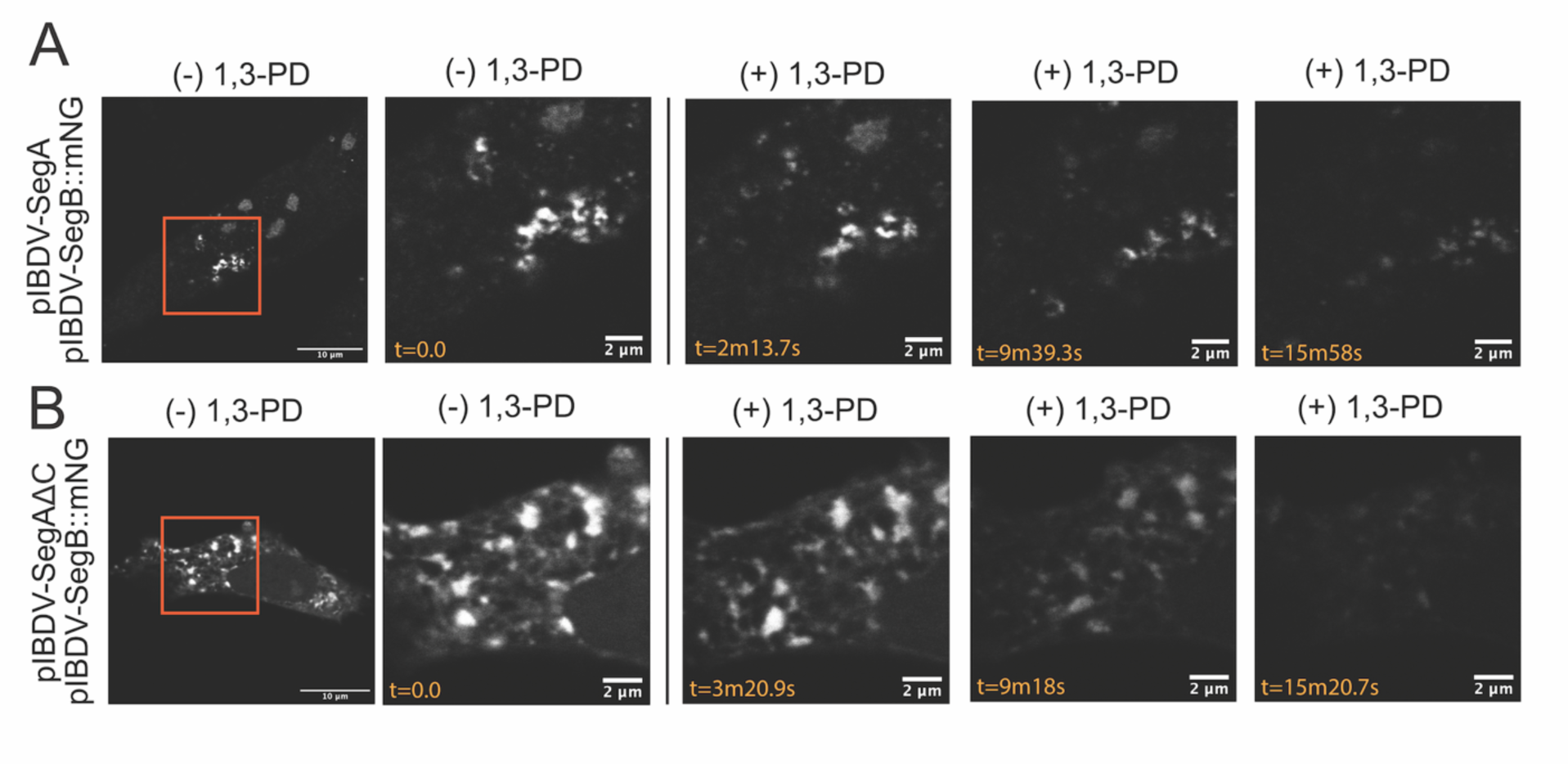
Puncta formed in the presence or absence of the VP3 C terminal IDR are sensitive to aliphatic diol treatment. DF-1 cells were transfected with a mixture of pIBDV-SegA and pIBDV-SegB::mNG (A), or DF-1 cells were transfected with a mixture of pIBDV-SegAΔC and pIBDV-SegB::mNG (B). Transfected cells were treated with 1,3-Propanediol (1,3-PD), while capturing time lapse images. Briefly, cells were initially maintained in media for acquisition, and at time (t)=0 seconds (s), 1,3-PD was added to achieve a final concentration of 4% (v/v). The leftmost images show whole cells (mNG signal shown in white), with stills from the inset regions (orange boxed regions) shown thereafter. The timestamp (t) of imaging is shown in seconds (s).

## Discussion

In previous work, we determined that the VFs of birnaviruses such as IBDV were formed through the process of LLPS (10, 33), but the molecular mechanism of how birnaviruses drive LLPS to form their VFs remained unknown. A major component of IBDV VFs is the VP3 protein (10, 18, 19). In this study, we used molecular dynamics simulations to robustly characterize the behavior of the predicted structure of the IBDV VP3 protein monomer and homodimer. We analyzed the predicted thermal motion (B-factor) and internal positional correlation of the atoms across the structure, and based on these data, we concluded that the 36 C-terminal amino acids of VP3 comprised a predicted highly dynamic IDR, a result which agreed with the predicted behavior of this region based on the protein sequence.

IDRs are frequently implicated as drivers of LLPS as their conformational plasticity is well-suited to the multivalent weak interactions that cause the phenomenon (34). We therefore hypothesized that VP3 acts as a scaffold for IBDV VFs, with its IDR as a key driver of LLPS. Consistent with this hypothesis, we discovered that the rate of cytoplasmic puncta formation was significantly reduced in the presence of VP3 lacking the C terminus, as compared to the presence of full-length wt VP3, with fewer punctate cells being observed and fewer puncta per cell. Moreover, the puncta that did form in the absence of the VP3 C terminus were significantly smaller, had a reduced mean fluorescence intensity, and were less “round” or “smooth” than those formed from wt VP3. Additionally, using FRAP, we discovered that the mobile fraction in puncta that formed in the absence of the VP3 C terminus was significantly lower (0.291) than the puncta that formed in the presence of full-length wt VP3 (0.704) (p<0.01), demonstrating that the VP3ΔC puncta were less liquid-like. Taken together, these data demonstrated that the C terminal IDR of VP3 was a potent driver of cytoplasmic puncta formation and influenced their liquidity.

However, three lines of evidence led us to hypothesize that rather than the VP3 C terminal IDR being the sole driver of LLPS, VP3 could be a member of a higher-order matrix that included other biomolecules to provide the required multivalency that drives LLPS: First, when expressed alone, the full-length wt VP3 protein did not spontaneously phase separate in DF-1 cells. This is in contrast to the scaffold protein of ReoV (µNS), which spontaneously phase separates in transfected cells in the absence of other viral components (15).Interestingly, VP3 has been reported to form small puncta in QM-7 cells stably expressing the protein (19). However, these puncta did not resemble the smooth or round structures that are characteristic of LLPS, and the authors did not assess their physical properties, so while there may be cell-dependent differences in the phenotype of VP3, the prevailing evidence is that VP3 does not phase separate alone. This suggests that the C terminal IDR of VP3 is not sufficient to induce LLPS. Second, removal of the VP3 C terminus did not completely abolish cytoplasmic puncta formation, suggesting that the VP3 C terminal IDR was not strictly required for puncta formation. Third, we observed that exogenous VP3 could be recruited to IBDV VFs in infected cells, even in the absence of the C terminus. Such behavior precluded the C-terminus being a requisite for VF compatibility. Moreover, as VP3 is known to dimerize, it is possible that when VP3ΔC was recruited to the VFs of infected cells, it may have formed heterodimers with full-length VP3 provided by the virus. LLPS requires multivalent intermolecular contacts, frequently three or more (34), and although the two protruding C-termini of the dimer were appealing candidates as contact sites to ensure multivalency associated with LLPS, the ability of the VP3ΔC to be recruited to VFs in infected cells suggested that deletion of at least one C-terminus did not remove this required valency. Therefore, instead of driving the initial formation of the cytoplasmic puncta, we hypothesize that the VP3 C-terminal IDR might interface with client biomolecules, thus influencing the physical properties of the resultant structures, for example by affecting the kinetics of the recruitment of biomolecules to VFs, or the packing and diffusive environment within the structures. In other words, the VP3 C terminal IDR could be important in modulating how liquid the VFs are.

Interestingly, we observed minimal differences in the dynamic behaviors of the puncta formed in the presence of full length wt VP3 and in the presence of VP3ΔC: both were freely mobile in the cytoplasm and appeared to undergo fusion events, and both were sensitive to treatment with small aliphatic diols, which are means of designating LLPS (31). It remains unknown why some properties of the puncta were more significantly altered by the removal of the VP3 C terminal IDR than others, however, as these live cell imaging assays are frequently applied qualitatively, it was not possible to draw quantitative conclusions as to how the presence or absence of the VP3 C terminus affected these behaviors. In addition, as the aliphatic diols broadly perturb the hydrogen bonding environment of the entire cell, they caused a notable toxicity in DF-1 cells, even at low concentrations. To mitigate cytotoxicity, we treated cells with 1,3-PD, a less-toxic compound in the same class as the more commonly used 1,6-HD, however, responses varied too widely between cells and between puncta within a single cell to quantitatively distinguish differences in the behavior of puncta formed in the presence of the full-length VP3 or VP3ΔC.

The most quantitative assay that measured the physical properties of the puncta was FRAP, and as mentioned above, these data were consistent with the VP3 C terminus promoting the liquidity of the puncta, as the mobile fraction measured for puncta formed in the presence of the full-length VP3 was significantly higher than in the puncta formed by VP3ΔC. Interestingly, we also observed that the recovery of the VP3ΔC puncta exhibited a shorter recovery half-life (0.341 seconds), whereas the recovery of the wt VP3 puncta was longer (3.517 seconds). Although both were very rapid in the context of FRAP, a faster recovery half-life did not intuitively pair with a substantially reduced mobile fraction, as rapid recovery is suggestive of liquidity, whereas a reduced mobile fraction is suggestive of rigidity. However, it is important to note that in our system, the mNG fluorophore was fused to the client protein VP1, rather than the scaffold protein VP3. Previous studies have indicated that VP3 directly binds VP1 via the 10 C-terminal amino acids of VP3 (35). As these are absent in the ΔC system, it is therefore possible that the unexpectedly rapid recovery half-life in the VP3ΔC puncta was due to the presence of a small amount of unbound VP1-mNG that was able to move more readily as it was not tethered to VP3, even if the rest of the puncta was more rigid. However, further work is needed to confirm this effect.

Taken together, our data demonstrate that the 36 C-terminal amino acids of IBDV VP3 comprised a predicted IDR that promoted the formation of cytoplasmic puncta and modulated their physical properties. We propose that VP3 interacts with other biomolecules to form a higher-order matrix that collectively drive LLPS in forming IBDV VFs, and that the VP3 C terminal IDR is important in modulating the liquidity of the structures. This is somewhat analogous to rotavirus NSP2, where the flexibility of the C terminal region (CTR) supports LLPS and the biogenesis of rotavirus VFs (17). Moreover, a single point mutation (K294E) in the NSP2 CTR was predicted to reduce its flexibility: Compared to wild-type NSP2, this mutation induced smaller and more numerous viroplasms in infected cells, severely impacted viral replication, and adversely affected condensation behavior with NSP5 *in-vitro (17)*. These data demonstrate the sensitive dependence of rotavirus viroplasm condensation dynamics and properties on the flexibility of the NSP2 CTR and showed that even small changes to critical protein regions can induce measurable effects on biomolecular condensates that directly impact viral replication.

Our data indicate that the C-terminus of IBDV VP3 is likewise an important LLPS-modulating region, and we are currently undertaking studies to identify if critical features and residues can act as determinants of VF properties. Furthermore, VP3 is known to bind phosphatidylinositol-3-phosphate (PI3P) on the cytoplasmic surface of early endosomes (EEs) and form a non-uniform coating (19). If VP3 acts as a modulator of LLPS, these regions may act as “nucleation sites” for more efficient VF formation. However, additional work on the interaction of LLPS-structures with cellular membranes is needed to make further conclusions.

## Materials and Methods

### AlphaFold

Structural modeling with AlphaFold2 was accomplished using the AlphaFold2 Colab notebook published by Deepmind (https://colab.research.google.com/github/deepmind/alphafold/blob/main/notebooks/AlphaF old.ipynb?pli=1), based on the amino acid sequence of the VP3 protein from IBDV strain PBG98, obtained from GenBank Accession <u>MT010364</u>. Processing was performed using the monomer model, with relaxation performed in GPU mode. Structural modeling with AlphaFold3 was accomplished using the AlphaFold Server (https://www.alphafoldserver.com). Structures were visualized with UCSF ChimeraX v.1.17.1 (UCSF, CA) (36).

### Predicted Aligned Error (PAE)

AlphaFold’s predicted error in the relative position of each pair of residues, measured in Angstroms, was extracted from the .zip archives generated by the AlphaFold Colab notebook by ingesting the “predicted_aligned_error.json” file with Python’s JSON module, and values were plotted as a heatmap with Plotly (Plotly Technologies Inc.), see supplemental material for more information.

### IUPred3 Disorder Plot

IUPred3 was accessed through the Eötvös Loránd University web portal (https://iupred3.elte.hu/) and run in short disorder mode with medium smoothing. Results were downloaded in JSON format and loaded using Python’s JSON module, and plotted as a line plot with Plotly.

### Sequence Entropy

The sequence entropy of VP3 was determined and plotted as a line plot with Plotly (see supplemental material for more information).

### Molecular dynamics (MD) simulations

To run the MD simulations and generate movies, the software GROMACS was installed on a fresh installation of Ubuntu 22.04, running on an AMD 3800X processor with 32GB of RAM. The GROMACS binary was compiled with the (-DGMX_GPU = ON) flag, with CUDA cores provided by an Nvidia GTX 1080Ti graphics coprocessor (Nvidia, CA). Each simulation was 20 nanoseconds in duration and performed using the OPLS-AA/L all-atom force field with a 2 femtosecond (fs) timestep. Energy minimization and equilibration were carried out before each simulation. The trajectories of every alpha carbon atom in VP3 were determined, and the absolute point positions were extracted from the trajectory data. Periodic boundary condition effects were corrected algorithmically (see supplemental material for more information), and the alpha carbons of VP3 were visualized as animated three-dimensional scatter plots of the atomic co-ordinates.

### B factor calculations

The B factor was calculated from the MD simulations (see supplemental material for more information), and the mean and standard error of the mean (SEM) plotted as a line plot with Plotly.

### Atomic motion correlation analysis

For each MD simulation, the Pairwise Pearson’s product-moment correlation coefficients were calculated for the fluctuations of all alpha carbons about their mean positions (see supplemental material for more information).

### Cell culture

The chicken DF-1 fibroblast cell-line (37) was obtained from the American Type Culture Collection (ATCC) (VA, USA). Cells were maintained in Dulbecco’s modified Eagle’s medium (DMEM) (ThermoFisher Scientific, MA, USA) supplemented with 10% heat-inactivated fetal bovine serum (FBS) (Gibco, ThermoFisher Scientific, MA, USA) and incubated in an atmosphere of 37℃ and 5% CO_2_.

### Plasmids

Plasmids expressing the following were used in this study: IBDV VP3 (untagged), IBDV VP3 fused to mNG, TC, or eGFP reporters at the N-terminus (mNG::VP3, TC::VP3, eGFP::VP3), IBDV VP3 lacking the C terminus fused to mNG at the N-terminus (mNG::VP3ΔC), IBDV reverse genetics (RG) plasmid encoding segment A (pIBDV-SegA) (33), IBDV RG plasmid encoding segment B (pIBDV-SegB) (33), Segment A lacking the C terminus (pIBDV-SegAΔC), SegmentB with an HA tag inserted between the 5’UTR and the coding region of VP1 (pIBDV-HA::SegB), and SegmentB with mNG fused at the C-terminus (pIBDV-SegB::mNG). Each of the inserts was cloned into the pSF-CAG-KAN vector (Sigma-Aldrich, DE), and the IBDV sequences of Segment A, B, and VP3 were designed based on the cell-culture adapted strain, PBG98 (33) (GenBank accession numbers <u>MT010364</u> and <u>MT010365</u>).

### Molecular Cloning

The vector and insert DNA were digested with appropriate restriction nucleases (New England Biolabs (NEB) (MA, USA)), in accordance with manufacturer protocols for each enzyme. The reactions were then separated by gel electrophoresis and the Monarch Gel Extraction kit (NEB) was utilized to extract DNA from gel slices. The DNA was quantified by nanodrop and the backbone and insert DNA were ligated with T4 DNA ligase (ThermoFisher Scientific), in accordance with manufacturer protocols, at 16℃ overnight. The ligation product was then used to transform chemically competent E-coli (NEB), according to manufacturer’s protocols. After transformation and outgrowth in SOC media (ThermoFisher Scientific), the mixture was streaked onto agar plates with the appropriate antibiotic selection marker (ThermoFisher Scientific). Plates were incubated at 37℃ overnight to allow colonies to develop. Colonies were selected and grown overnight in LB broth (ThermoFisher Scientific), in the presence of selection antibiotic, at 37℃ with shaking at 200rpm. A 1mL aliquot of the broth was reserved from each of these outgrowths, and the remainder was processed using the Spin miniprep kit (Qiagen, MD), and screened by an analytical restriction digest. Once a positive sample was identified, additional LB broth supplemented with the appropriate antibiotic selection was inoculated with the 1mL reserved aliquot of the positive outgrowth and incubated overnight at 37℃ with shaking at 200rpm and processed using the Spin Maxiprep kit (Qiagen), with final elution in molecular biology grade water. The DNA was quantified by nanodrop, diluted with molecular biology grade water to 1,000ng/µL, and stored at -20℃.

### Transfection

DF-1 cells were transfected with the relevant plasmids using Lipofectamine 2000 transfection reagent (ThermoFisher Scientific), according to the manufacturer’s recommendations for the size of the culture vessel. Briefly, the lipofectamine 2000 reagent and DNA were diluted in Opti-MEM media (Gibco, ThermoFisher Scientific), mixed, and incubated for 20 minutes at room temperature (RT) before application to adherent cells dropwise into the media.

### Virus

A molecular clone of the cell-culture adapted IBDV strain PBG98 was rescued by transfecting DF-1 cells with the RG plasmids SegmentA and SegmentB, and monitoring cells for cytopathic effect (cpe) as previously described (33). The resulting virus was titrated in DF-1 cells and the titer quantified by the tissue culture infectious dose-50 (TCID_50_) method described by Reed and Muench (38). Unless otherwise stated, DF-1 cells were infected at a multiplicity of infection (MOI) of 5 in downstream experiments.

### Immunofluorescence microscopy

12-well tissue culture treated plates (Corning, AZ) were populated with 18mm No.1 glass cover slips and seeded with DF-1 cells and maintained as described (see cell culture). Cells were washed with sterile phosphate-buffered saline (PBS), fixed in 4% (v/v) paraformaldehyde (PFA) in PBS for fifteen minutes at RT, and washed again with PBS. Cells were then permeabilized with 0.1% (v/v) Triton-X 100 in PBS for 15 minutes, and incubated with a blocking buffer of 4% (w/v) of bovine serum albumin (BSA) in PBS for one hour at RT. The primary antibody was then diluted in blocking buffer, and cells were incubated in this solution at RT for one hour with rocking. After incubation, the primary antibody solution was aspirated and the cells were washed with PBS. The secondary antibody conjugated to a fluorophore was then diluted in blocking buffer, and cells were incubated in this solution for one hour at RT with rocking. Cells were washed with PBS, counterstained with DAPI and cover slips were mounted onto slides prior to imaging.

### Puncta quantification and measurement of morphology

Puncta were quantified by acquiring mosaic images with a Leica Stellaris 8 scanning confocal microscope. Cells were selected at random from the fields and puncta were counted manually. Fifty cells were analyzed for each of 3 independent trials for each condition. Measurement of puncta morphology in cells was accomplished by automatic segmentation. Briefly, images of 50 single cells were captured with a Zeiss LSM980 Airyscan scanning confocal microscope and processed using a custom Python script which performed watershed segmentation to isolate puncta. Segmentation was manually reviewed prior to measurement. The script is available on Github (https://github.com/LadInTheLab/vp3-llps), and the algorithm utilizes the Scikit-image library for segmentation and measurement. Statistics were calculated by grouping puncta by condition (wt or ΔC) and, after a Shapiro-Wilk test determined that the assumption of normality was not valid, a Mann-Whitney U Test was performed between the groups for each metric.

### Live cell imaging

DF-1 cells were seeded in confocal dishes with 20mm No.1 glass bottoms (MatTek, MA) at a density of 2 x 10^!^ cells per dish. Cells were imaged in a live-cell imaging chamber of a Zeiss LSM 980 laser scanning confocal microscope and maintained in a 5% CO2 environment at 37 Celsius during imaging. The microscope was configured to minimize laser power (and thus photobleaching) without introducing excess noise, and with Z-drift compensation configured to run every 5 frames to maintain a constant imaging plane.

### Aliphatic Diol Treatment

DF-1 cells were seeded in glass-bottom confocal dishes with 20mm No.1 glass bottoms at a density of 2 x 10^!^ cells per dish. Immediately prior to imaging, the media was removed from the cells and 100 µl of fresh media added. Cells were imaged with a Zeiss LSM 980 laser scanning confocal microscope configured to minimize laser power (and thus photobleaching) without introducing excessive noise, and with Z-drift compensation configured to run every 5 frames. An open-ended time-lapse acquisition was then configured. Thirty seconds after acquisition began, 900µl of media containing 4.44% (v/v) of the specified aliphatic diol (Beantown Chemical, NH) was carefully added to the dish by pipette, for a final concentration of 4% (v/v), with continual imaging. Image acquisition was continued until photobleaching, cell drift or toxicity became excessive.

### FRAP

FRAP was performed using a Zeiss LSM 980 laser scanning confocal microscope. The imaging area was cropped to maximize the proportion of the scanned area occupied by the puncta of interest, and a circular measurement region of interest (ROI) was established, centered about a point ROI. The system was configured to capture a 256x256 pixel image every 75.95 milliseconds with a 1AU pinhole. The experiment was configured to capture 5 pre-bleach frames, followed by a single-frame 488nm bleaching pulse at 10% power at the point ROI, followed by continuous acquisition of frames thereafter. Extraction of intensity data was performed in Fiji (v2.3.0/1.53f, open source). Unmodified files from the Zeiss microscope (.czi files) were loaded with BioFormats importer, and 3 ROIs were established on the images: “Bleach”, an ROI encompassing the region bleached in the first frame after the bleaching pulse, “puncta”, an ROI encompassing the entire puncta, and “background”, an ROI outside the puncta. The multi-measure function was then used to capture mean intensity data for each ROI across every frame of the experiment, the results of which were exported as comma-separated value (.csv) files for analysis. This analysis was performed in Python, with a script available at (https://github.com/LadInTheLab/vp3-llps). The script was based on easyFRAP-web (32). For more information, please see supplemental materials.

### Statistics and Plotting

Statistics and plotting were performed in Python unless otherwise specified. Statistical calculations were performed using the SciPy package (39), and plotted using the Plotly package (40).

## Acknowledgements.

This work was supported by University of Maryland startup funds to A. J. Broadbent. In addition, we acknowledge the Imaging Core Facility in the department of Cell Biology and Molecular Genetics at the University of Maryland, College Park for use of the Zeiss LSM980 Airyscan and Leica Stellaris 8. Purchase of the Zeiss LSM 980 Airyscan 2 was supported by Award Number 1S10OD025223-01A1 from the National Institute of Health (NIH). Purchase of the Leica Stellaris 8 was supported by Award Number 1S10ODO34260 from the NIH. The funders had no role in study design, data collection and interpretation, or the decision to submit the work for publication.

